# Vasoconstriction in isolated goat aorta does not increase mean aortic pressure

**DOI:** 10.1101/2021.01.30.428980

**Authors:** Naveen Gangadharan, V Aravindhan, Benjamin Jebaraj, Shikha Mary Zachariah, Suresh Devasahayam, G Saravana Kumar, Sathya Subramani

**Affiliations:** Department of Engineering Design, Indian Institute of Technology Madras, Tamil Nadu, India; Department of Physiology, Christian Medical College Vellore, 632002, Tamil Nadu, India; Department of Bioengineering, Christian Medical College Vellore, 632002, Tamil Nadu, India

**Keywords:** aorta, small artery, resistance, compliance, adrenaline, vasoconstriction, mean arterial pressure

## Abstract

Vasoconstriction in small arteries and arterioles is known to increase resistance to flow, while vasoconstriction in large arteries and aorta is known to decrease their compliance. Besides this general understanding, there is no systematic documentation on what happens to small artery compliance and large artery resistance during vasoconstriction and the corresponding alterations in vascular pressure. The aim of the study is to assess the effect of adrenaline on goat aortae and small arteries in terms of resistance and compliance.

Isolated goat aortae and small arteries were perfused with a pulsatile pump and lumen pressure was recorded before and after addition of adrenaline. In the aortae, systolic pressure increased, diastolic pressure decreased, pulse pressure increased (p = 0.018, WSR); but the mean pressure remained the same (p = 0.357, WSR). Small artery vasoconstriction caused an increase in systolic, diastolic and mean pressures (p = 0.028, WSR). Using length, radius, and thickness data from the tissues and the tubes of the experimental set-up, electrical models were simulated to understand the biological data. The simulations allow us to infer that vasoconstriction in aorta leads to a reduction in compliance, but an increase in resistance if any, is not sufficient to change the mean aortic pressure. On the other hand, vasoconstriction in small arteries increases resistance, but a decrease in compliance if any, does not affect any of the four pressure parameters measured. Vasoconstriction in aorta decreases compliance and therefore increases pulse pressure but does not change resistance significantly enough to alter mean pressure.

**Key Points Summary:**

- The main aim of the study is to understand where exactly resistance (*R*) and compliance (*C*) components of the vasculature occur. There is no definitive evidence for the effect of large artery vasoconstriction on resistance and hence the mean arterial pressure.
- The manuscript presents biological experiments studying the pressure response of goat aorta and small arteries to adrenaline (*invitro*) and the interpretations using equivalent electrical models.
- The study shows that in aorta and large arteries, vasoconstriction does not lead to a reduction in lumen diameter sufficient to cause a rise in resistance and mean pressure as compared to small arteries.
- Knowledge of exact location of *R* and *C* in the arterial tree enables re-assessment of the differential action of vasoactive drugs on resistance versus compliance vessels once we resolve beat-to-beat *R* and *C* changes in response to a drug. This way antihypertensive therapy can be tailored to address the specific cause of the type of hypertension.

## Introduction

Arterial compliance and resistance are important determinants of arterial blood pressure besides cardiac performance. Compliance is a measure of the ease with which the vessel can expand when a volume of blood is added to it. Technically, compliance is the change in the volume of the vessel (the container), per unit change in pressure in the vessel^1^. The elasticity of the elastic fibers in the vessel wall, as well as the tone of smooth muscles there determine the arterial compliance^2,3^. The same components also confer the property of elastic recoil to the artery, which can be thought of as the ease with which the vessel elastically recoils to a smaller volume, when blood runs off from the vessel. When systolic and diastolic pressures are considered independently, systolic pressure is inversely related to arterial compliance while diastolic pressure varies directly with compliance or rather, inversely with arterial elastance (the inverse of compliance)^4^.

The reason for calling the aorta and large arteries as Windkessel vessels is that they have some degree of compliance/elastance that allows them to minimize pulse pressure by limiting the systolic pressure from rising too high and diastolic pressure from falling too low^5,6^. Loss of compliance/inverse elastance as a person ages can lead to isolated systolic hypertension (ISH), where the systolic pressure is more than 140 mmHg while the diastolic pressure is less than the cut-off value used to define essential hypertension, i.e., less than 90 mmHg^7^. Such age-related loss of compliance is generally attributed to loss of elasticity due to degradation of elastic fibres in the vessel wall. Loss of elasticity is a slow aging process, occurring over years^8,9,10^. However, aorta has considerable smooth muscle in their tunica media, interspersed with which the elastic tissue is found^11^. The role of smooth muscle present in the walls of aorta and large arteries is less appreciated^12^. Vasoconstriction of aorta should lead to an acute reduction in its compliance and this component of compliance loss must be reversible with vasodilator drugs. That must be the rationale for treating ISH with vasodilator drugs^13,14^. However, it is not clear if vasoconstriction in aorta has an effect on aortic resistance. While it is well known that arteriolar resistance increases with vasoconstrictors, it is not clear if aortic (and large artery) resistance, while not being as significant as compared to arteriolar resistance, increases due to vasoconstriction^15,16^. This would enable targeted anti-hypertensive therapy based on the differential action of a vasoactive drug on resistance versus compliance vessels. One aim of this study is to assess the effect of adrenaline, a known vasoconstrictor on aortic resistance and compliance as compared to small arteries. Aortic smooth muscle, both circular and longitudinal, contract in response to adrenergic stimulation while in the arterioles the circular muscles contract and longitudinal muscles relax^11^. The questions we asked were whether the contractile response of the smooth muscle in the walls of aorta (and large arteries) leads to (1) an acute change in compliance, or (2) a change in resistance, or (3) both.

Vasoconstriction in the walls of aorta and the large arteries will lead to a decrease in compliance of the vessel walls and/or reduction in vessel circumference. The decreased compliance will lead to the following changes in the pressure pulse when perfused with a pulsatile flow (by a peristaltic pump): increase in peak (systolic) pressure and decrease in trough (diastolic) pressure, with a consequent increase in pulse pressure. The mean pressure during the pulse however is not expected to change with only changing compliance. If such aortic/arterial vasoconstriction increases resistance, then the peak, trough and mean pressures will all increase. Our assay therefore involved measurement of lumen pressure with a transducer connected to a cannula placed in an isolated segment of aorta.

The research question can be better understood when considered in relation to vasoconstriction in arterioles and small arteries. While the aorta and large arteries are classified as Windkessel vessels, the small arteries and arterioles being the major region of pressure drop in circulation are referred to as resistance vessels^17^. The resistance vessels are known to explicitly alter their resistance by altering their lumen diameter in the presence of vasomodulators^18^. If the resistance vessels are perfused with a pulsatile pump, a pulsatile pressure, with peaks and troughs similar to physiological systolic and diastolic pressures just like the aorta will be recorded. If the resistance increases due to vasoconstriction, then mean pressure, peak and trough pressures will increase. While this is the expected response in arterioles and small arteries, the response of aorta to vasoconstrictors is not clear. Our aim here is to compare the responses of aorta and small arteries to adrenaline, a known vasoconstrictor, and draw conclusions about their resistance and compliance.

## Methods

This study was approved by the Institutional Review Board of Christian medical College, Vellore, IRB no: 9930, 17/02/2016.

Intravascular pressure was measured in isolated lengths of goat aortae and small arteries perfused with mammalian ECF solution with a pulsatile pump (delivering a constant stroke volume), before and after administration of adrenaline. The radius and thickness of the vessel segment studied was measured using a Vernier calipers before the intervention without squeezing the vessel.

### Isolation of Aorta

Fresh goat heart (*Capra hircus*) with a considerable length of thoracic aorta was procured from a registered slaughterhouse (Rafi’s slaughterhouse, accreditation number: 309/13, Co-ordinates 12.93°N 79.13°E) and transported to the lab in ice cold extra-cellular (EC) solution on the day of the experiment. The heart was washed well in cold EC solution. The ascending aorta was identified after removing the pericardial sac and fat. A cylindrical segment of about 70 mm length was dissected from its origin to the point before it branches. The dissected segment was then transferred to a petri dish containing cold EC solution, following which the adventitious tissues were removed. It was ensured that the segment was free of branches and if there were branches, they were ligated.

### Experiment set-up for flow experiments

Two metal cannulae were placed on either sides of the aortic cylinder and secured with ties. They served as inlet and outlet of the aortic cylinder. The inlet was connected to a pulsatile pump (Harvard Apparatus, rodent blood pump model 1407) through a length of non-compliant plastic tubing and the outlet was connected to similar tubing which was left to drain freely. Extracellular solution was perfused through the aortic cylinder at 37°C. For tissue viability, the EC solution was bubbled with carbogen (95% O_2_ and 5% CO_2_). Stroke rate of the pump was set at 80 strokes/min and the flow was calibrated to 1.33 ml/stroke.

A cannula was then inserted into the aortic segment, facing the flow at the inlet side and a commercially available pressure transducer was connected to the intra-aortic cannula to record fluid pressure at the inlet of the segment. Pressure waveform was recorded in a validated data acquisition system (CMCdaq) at 1000Hz sampling rate. Pressure was recorded under control conditions and when the recordings were stable, adrenaline was added (10μmol/L) to the perfusate. Pressure was recorded over 15-20 min after adrenaline addition. The experiment set up is shown in Figure 1.

**Figure 1:**
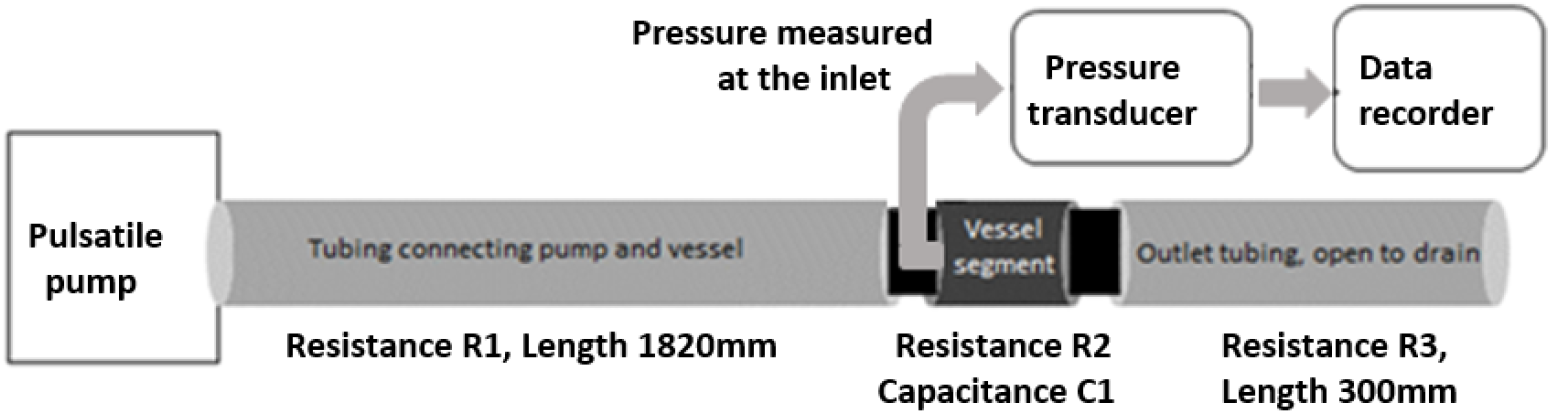
Schematic of the experimental set up. The “vessel segment” is either a section of aortic cylinder or small artery cylinder

### Isolation of small artery

Fresh goat legs (*Capra hircus*) were procured from a registered slaughterhouse (*ibid*). A small artery was identified from a vascular bundle close to the muscle below the knee by patency of its lumen. The desired section of the artery was dissected and transferred to cold EC solution. After removing the adventitious tissue attached to the artery, it was cut into segments of 30 - 35 mm length, without any side branches. The small artery segment was cannulated and perfused in the same manner as the aortic segment. Flow rate was calibrated to a smaller value (0.1 ml/stroke).

### Solutions Used

The mammalian ECF solution used in the experiments had the following composition (in mmol/L): KCl 3; NaCl 100; CaCl_2_ 1.3; Na_2_HPO_4_ 2; NaHCO_3_ 25; NaH_2_PO_4_ 0.5; MgCl_2_ 2; HEPES 10; Glucose 5; pH of the solution was corrected to 7.4 with 1 mol/L NaOH; the salts were purchased from SIGMA. Adrenaline bitartrate was purchased from a certified pharmacy. 10 μmol/L concentration of Adrenaline was used for the experiments.

### Electrical Analogue of the experimental set up (used for simulations)

An electrical analogue of the experimental set up shown in Figure 1 was devised to simulate variations in resistance and capacitance across the whole functional spectrum and assess what the changes in voltage (equivalent to pressure) would be. The experimental set up shown in Figure 1 consists of a time varying flow pump, a resistive connecting tube, compliant blood vessel followed by a draining tube. Resistance to fluid flow (*R*) is the ratio of pressure drop to flow in a vessel. It depends on the dimensions of the vessel (cross-sectional area and length) and the viscosity of the fluid. In the case of a cylindrical vessel of cross-sectional diameter *d* or radius *r*, cross-sectional area *A*, length *l*, carrying a fluid with viscosity *η,* the resistance to flow can be calculated as:

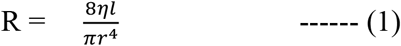

The capacitance of a vessel under time-varying flow is the ratio of the change in pressure to change in volume. It depends on the dimensions of the vessel and the elastic compliance of the vessel wall. The capacitance can be calculated as:

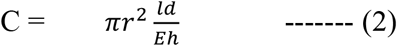

Here ‘E’ is the elastic modulus or Young’s modulus, and ‘h’, the thickness of the vessel. The SI unit for resistance to fluid flow is Pa·m^3^·s and for capacitance it is Pa^-1^·m^3^, although sometimes these are designated Ohm and Farad in keeping with electrical notation for analogous electrical circuit calculation. The SI unit for Young’s modulus^5^ is Pa and that for viscosity is mPa. s. The experimental arrangement can now be represented by (a) an alternating voltage source to represent the pump (a time varying voltage source is used in the electrical equivalent circuit, as the time varying flow pump is equivalent to a time-varying pressure pump with a series resistance (using Thevenin equivalent from circuit theory) (b) a resistance R1 for the tube connecting the pump and the vessel, (c) a resistance and capacitance, R2 and C1 for the vessel segment, and (d) a resistance R3 for the tube that drains the fluid to the open atmospheric pressure or reference pressure. The following Figure 2 shows the equivalent electrical circuit.

**Figure 2:**
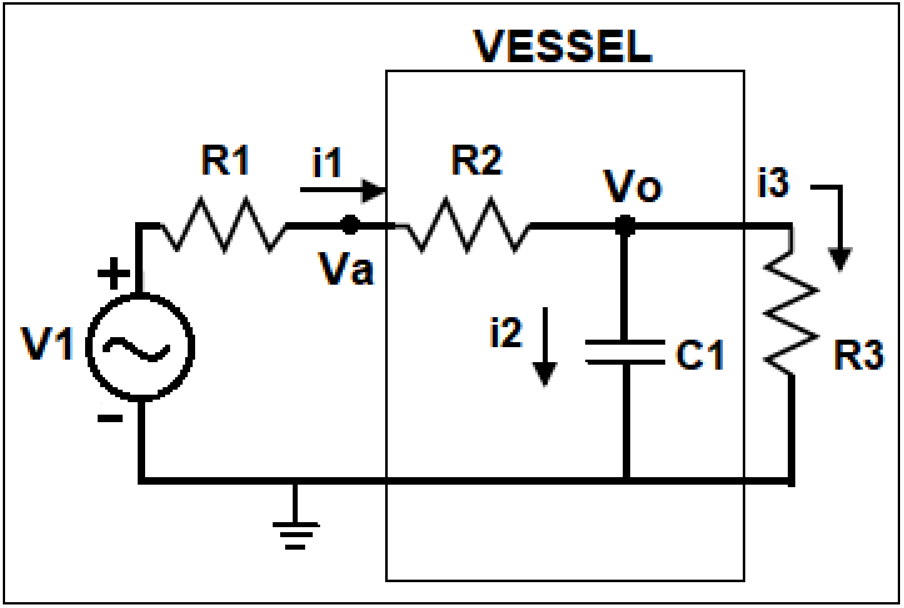
Electrical circuit equivalent of the experimental set up shown in Figure 1

This circuit is a linearized representation of the fluid flow in the experimental arrangement. To make the electrical simulations meaningful, it is necessary to set the initial values of R1, R2, R3 and C1 close to values representative of the components of the experimental set up (see Table 1). R2 and C1 should be close to resistance and capacitance of the aorta / small artery and R1 and R3 should be similar to the inlet and outlet resistances.

The initial values for the resistances R1 to R3 and compliance for the aorta and small artery were estimated from actual lengths, radii and thickness of the tubings and the vessels in the experimental set up using the equations (1) and (2). Viscosity of the perfusate was taken as 1 mPa.s. R1 and R3 were constant across all experiments. R2 and C1 were changed in the simulated electrical circuit, to study the pressure responses for entire ranges of R2 and C1 for the aorta and small artery separately. The pressure response to adrenaline in the biological samples was compared with the voltage response in the simulation to changing R2 and C1, and conclusions drawn as to whether adrenaline changed R2 or C1 or both in aorta versus small artery.

### Imaging experiments

Circular rings of about 1mm thickness were cut out from the ascending aorta and placed in a Petri-dish and perfused with extracellular solution at 37°C. A standardized camera system was used for imaging the vessel vertically from the top. The vessel behavior in the absence and presence of (10μmol/L) adrenaline was recorded using the camera as a continuous video. After addition of adrenaline, recording was continued for 15 minutes. Post-hoc analysis of the recording was performed to determine the lumen radius of the aortic ring per frame using a calibrated custom program written for diameter calculation. The program takes the video as input and generates a csv file of frame number versus diameter of the aortic ring per frame. The diameter can be mapped to the time of addition of Adrenaline based on recorded timestamps.

### Statistical analysis

SPSS software ver.16.0 was employed to do the statistical analyses. Comparison of intra-vascular pressures (peak, trough and mean) before and after addition of adrenaline in aorta experiments and small artery experiments was done using Wilcoxon’s signed rank (WSR) test. P value of 0.05 was considered statistically significant.

## Results

### Results of biological experiments

Effect of adrenaline on aortic pressure: Pressure pulses were recorded from the aortic segments. Raw tracing of the aortic pressure recording before and after adrenaline is shown in Figure 3A. Adrenaline increased peak pressure (3B), reduced trough pressure (3C), but did not change mean pressure (3D) in aortic segments (n = 7). It took nearly 15 minutes for the full response to adrenaline to develop. The changes in peak pressure, trough pressure and pulse pressure were statistically significant (p = 0.018, WSR) while there was no significant change in mean pressure (p = 0.357, WSR).

**Figure 3:**
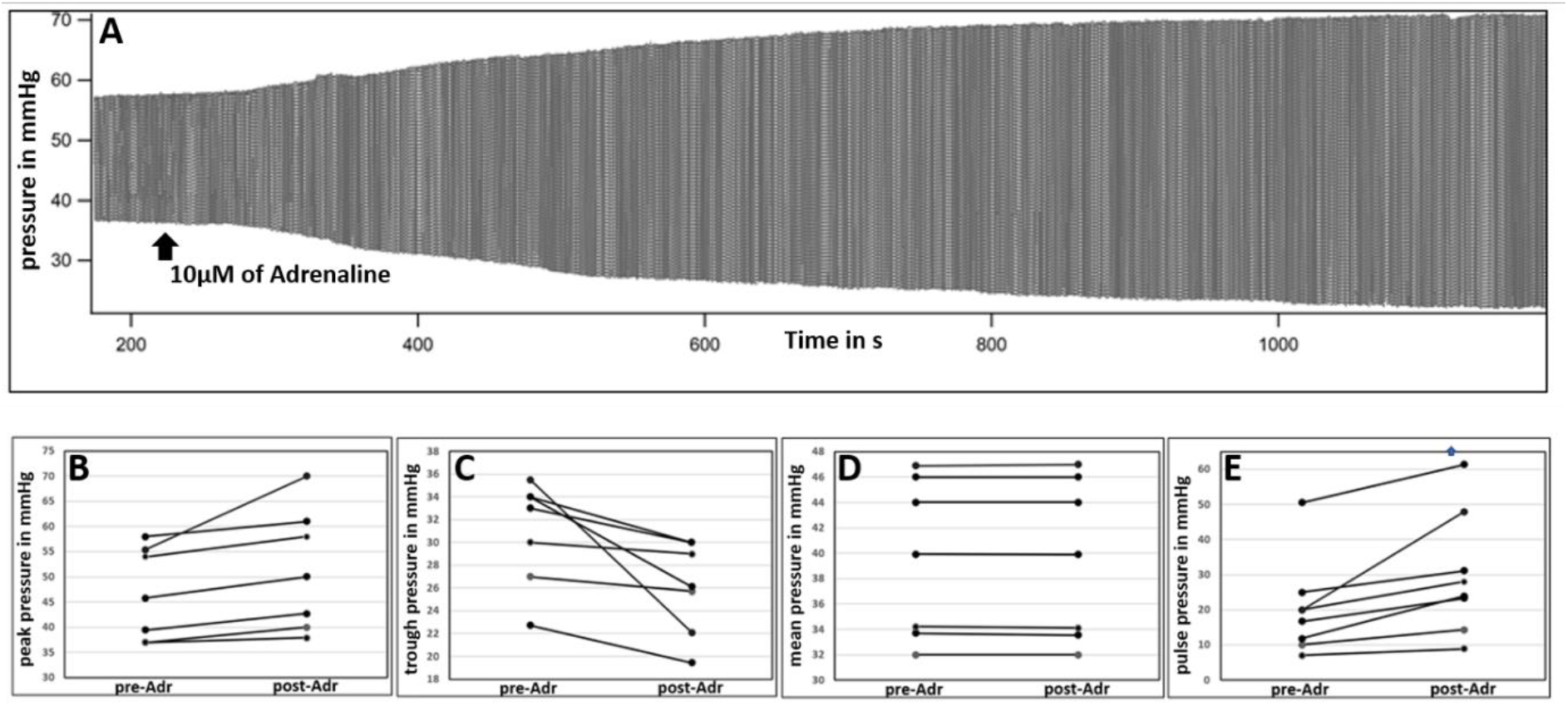
(A) Raw tracing of pressure in goat aortic cylinder; Scatter plot of (B) peak pressure (C) Trough pressure (D) Mean pressure (E) Pulse before and after addition of 10 μmol/L adrenaline in 7 samples of aorta

Effect of adrenaline on pressure in small arteries: To place the observations in aorta in context, we studied the effect of adrenaline on small arteries. Raw tracing of pressure recorded from a small artery before and after adrenaline is shown in Figure 4A. Adrenaline increased peak (4B), trough (4C), and mean vascular pressures (4D) in small arterial segments (n = 6). The full response to adrenaline was almost instantaneous in the small arteries. The rise in peak, trough and mean pressures were statistically significant (p = 0.028, WSR). The rise in pulse pressure was not statistically significant (p=0.075, WSR).

**Figure 4:**
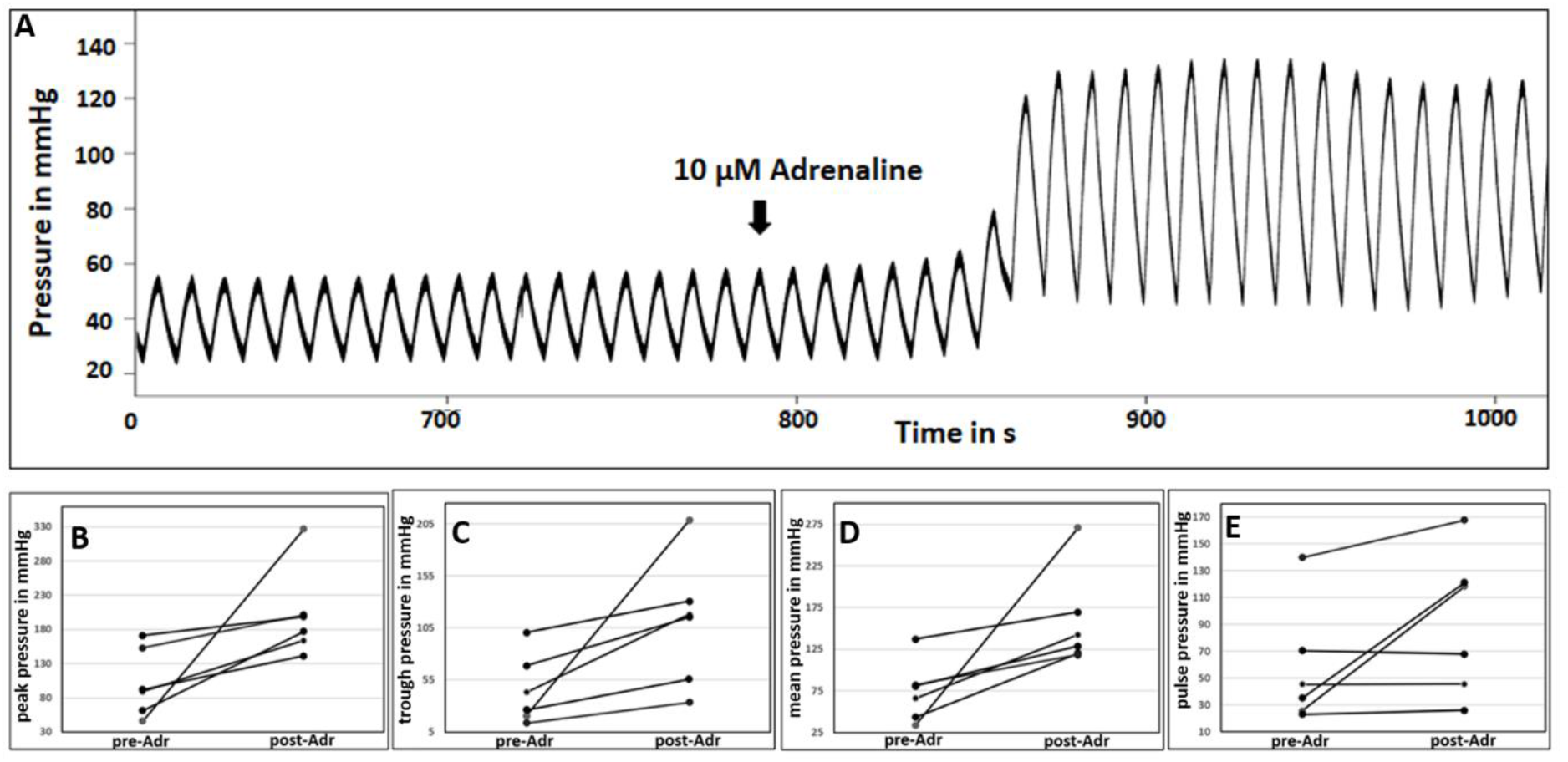
(A) Raw tracing of pressure in small artery cylinder; Scatter plot of (B) Peak pressure (C) Trough pressure (D) Mean pressure (E) Pulse before and after addition of 10 μmol/L adrenaline in 6 samples of small artery

### Results of simulations

Simulations were performed with an electrical circuit analogous to the biological experiment set up, to understand the pressure response of aorta and small artery to adrenaline. Initial values for the resistances R1-3, and C1 were calculated from measurements made in one sample each of aorta and small artery. The values of the calculated resistances and capacitances are shown in Table 1.

**Table 1:** Values used to calculate the resistances R1 – 3 and Capacitance C1 (refer Figures 1 and 2) for the electrical simulation

|  | length (l)<br>of the tube<br>or vessel | radius (r)<br>of the tube<br>or vessel | Resistance or<br>Capacitance for<br>fluid flow | Electrical<br>equivalents<br>used for the<br>simulation |
| --- | --- | --- | --- | --- |
| R1 (the non-compliant tube from the pump to aorta) | 1820 mm | 5 mm | $119 \times 10^6 \text{ Pa} \cdot \text{m}^3 \cdot \text{s}$ | $119 \times 10^6 \text{ ohms}$ |
| R2 of aorta- (for one sample) | 70 mm | 5 mm | $285.2 \times 10^3 \text{ Pa} \cdot \text{m}^3 \cdot \text{s}$ | $285.2 \times 10^3 \text{ ohms}$ |
| C1 of aorta<br>(thickness (h) of the vessel = 1mm) | 70mm | 5 mm | $1.1 \times 10^{-9} \text{ Pa}^{-1} \cdot \text{m}^3$ | $1.1 \times 10^{-9} \text{ farads}$ |
| R2 of small artery – (for one sample) | 31 mm | 0.25 mm | $20209 \times 10^6 \text{ Pa} \cdot \text{m}^3 \cdot \text{s}$ | $20209 \times 10^6 \text{ ohms}$ |
| C1 of small artery<br>(thickness (h) of the vessel = 0.25 mm) | 31 mm | 0.25 mm | $4.75 \times 10^{(-21)} \text{ Pa}^{-1} \cdot \text{m}^3$ | $4.75 \times 10^{(-21)} \text{ farads}$ |
| R3 of the non-compliant tube at the outlet end | 347mm | 1 mm | $884 \times 10^6 \text{ Pa} \cdot \text{m}^3 \cdot \text{s}$ | $884 \times 10^6 \text{ ohms}$ |

The electrical circuit equivalent (Figure 2) of the experimental set-up (Figure 1) was used to generate a sine voltage/current signal (equivalent of a pressure pulse or flow generated by heart). Voltage at the R1-R2 node (equivalent to recorded pressure in the vessel in biological experiments stated above) was studied with simulations where either R2 or C1 or both were altered. Values for R1 and R3 were constant for all simulations. For aorta simulations, R2 and C1 were set initially to the equivalents for aortic resistance and compliance (as in Table 1). The values of R2 and C1 were then independently varied to understand how they affect peak and trough pressure, pulse pressure and mean pressure.

Similarly, for small artery simulations, R2 and C1 were set at equivalents for arterial resistance and compliance. The voltage signal was set to correspond to the measured pressure in the tissue experiments.

#### Aorta simulations

##### Simulation 1

(Figure 5, top panels): At the arrow shown, R2 was changed to a higher value (equivalent to 50% reduction in radius from 5 mm to 2.5 mm) while C1 remained unchanged. R1 and R3 (of the non-compliant tubes) were chosen as in table 1. It was observed that peak, trough, mean and pulse pressures of the pressure pulse remained unchanged for such a gross reduction in radius.

**Figure 5:**
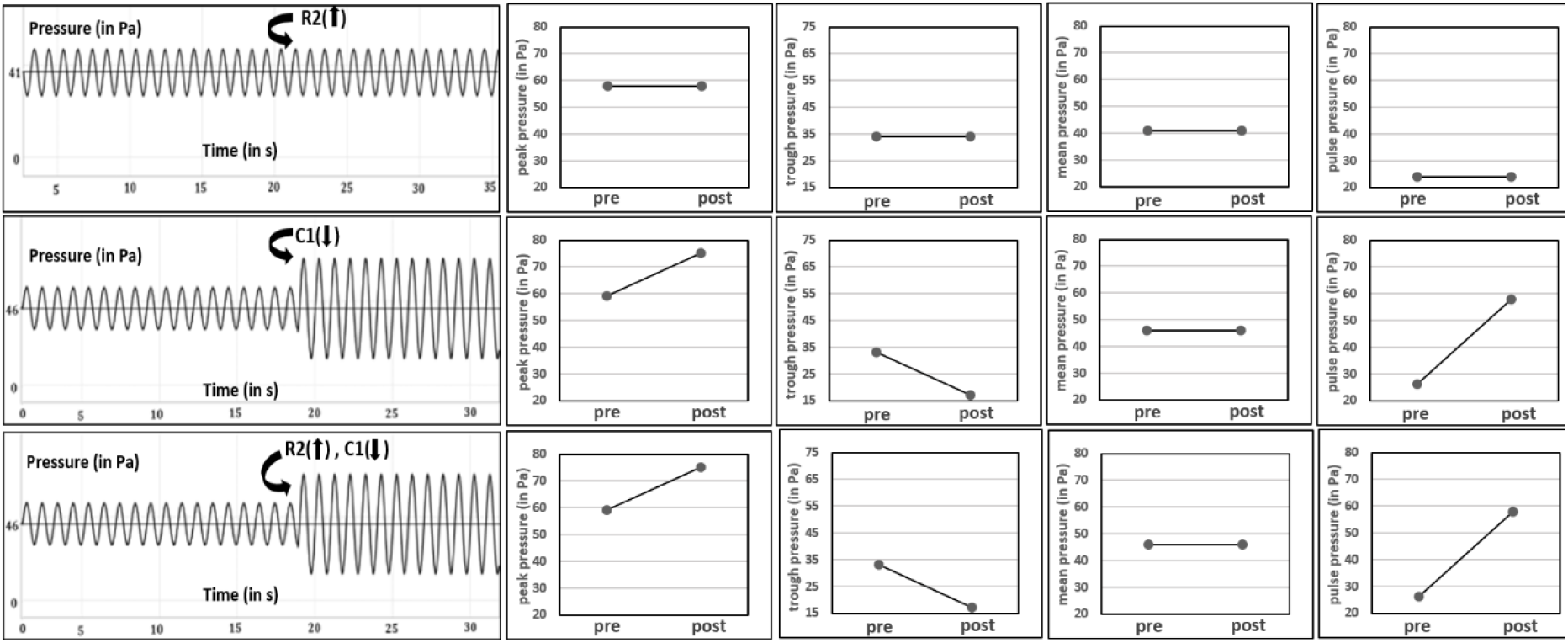
Simulations for aorta, where R2 was increased by 50% reduction in radius (top panels); C1 was decreased by 90%, (middle row of panels); R2 was increased and C1 decreased simultaneously (bottom panels). V1 was set as 34 Pa, 1 Hz sine, as observed in the biological experiments

##### Simulation 2

(Figure 5, middle row of panels): When C1 was reduced (90% reduction in compliance from 1.1x 10^-9^ to 0.11x 10^-9^ Pa^-1^·m^3^) and R2 left unchanged, there was an increase in peak pressure, decrease in trough pressure and there was no change in mean pressure.

##### Simulation 3

(Figure 5, bottom panels): When R2 was increased (50% reduction as in simulation 1) and C1 reduced simultaneously, (90% reduction in compliance as in simulation 2), it was seen that peak pressure increased and trough pressure dropped, while the mean pressure remained the same.

The results from the biological experiments with addition of adrenaline on aorta resemble simulations 2 and 3 in Figure 5. Responses in simulations 2 and 3 were identical. Since simulation 1 involving a change in resistance alone did not have any effect, it can be inferred that all of the observed change in simulation 3 was just due to changing compliance. This observation allows us to infer that adrenaline decreases compliance of aorta, while not changing its resistance sufficiently enough to see an increase in mean pressure.

The calculations as above were extended to predict pressure changes for:

(a) all possible increases in resistance starting from (285.2 x 10^3^ Pa·m^3^·s), which translates to reductions in radius starting from 5 mm (see, Figure 6, top panels); (b) all possible reductions in compliance, starting from 1.1x 10^-9^ Pa^-1^·m^3^ (see, Figure 6, bottom panels) if there was aortic vasoconstriction.

**Figure 6:**
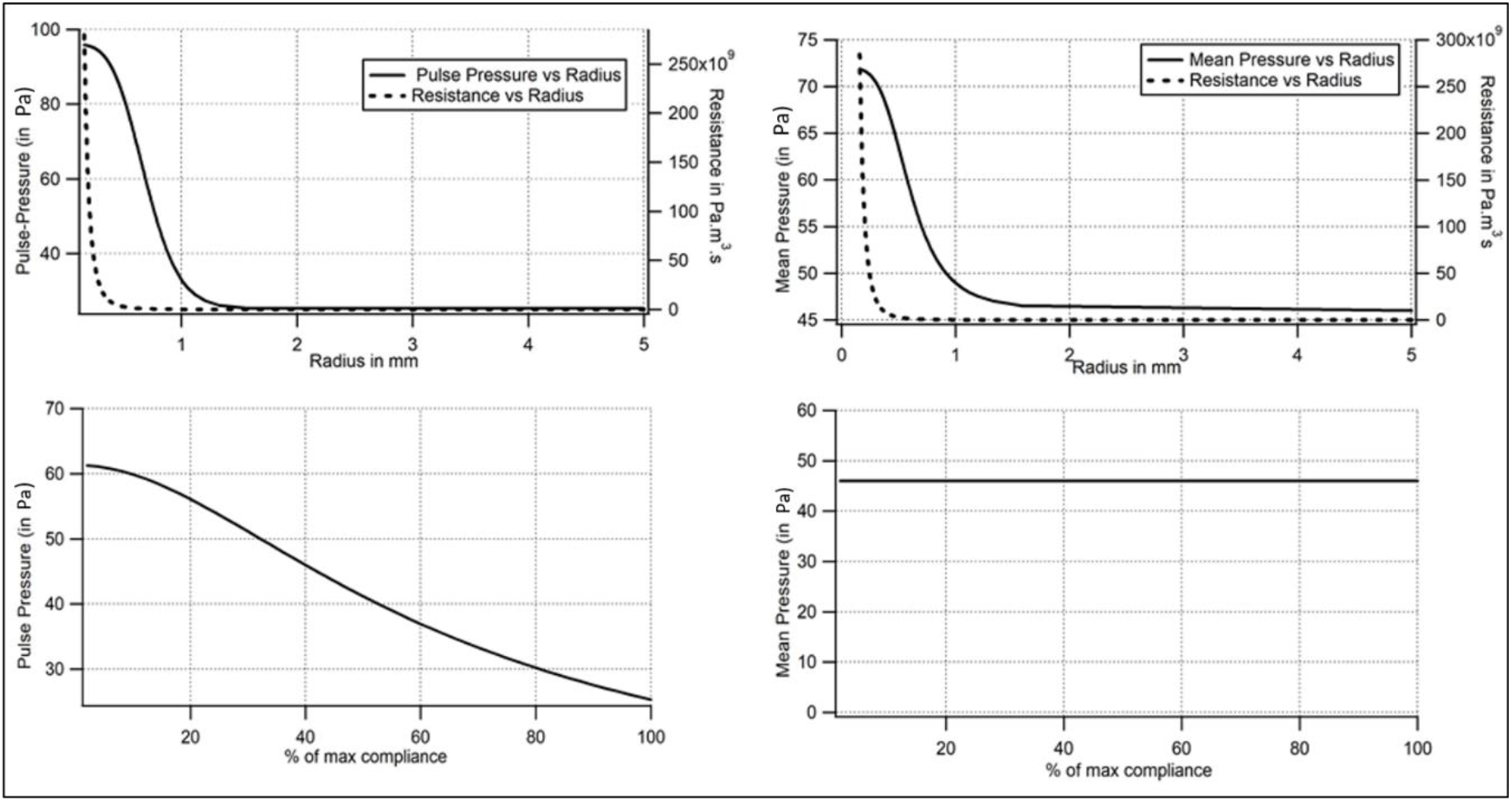
Analytical results of the effect of all possible reductions of radius on pulse pressure and mean pressure (top row of panels) in aorta simulations; analytical results of the effect of all possible reductions in compliance on pulse pressure and mean pressure (bottom panels) in aorta simulations.

Changes in mean pressure and pulse pressure were then plotted against actual radius (top panels) or percent reduction in capacitance (bottom panels). The axes of the figures are such that vasoconstriction is shown as a leftward change of X axis parameters.

It is seen that there is no significant change in mean pressure or pulse pressure till the radius reduces from 5 mm to about 2 mm. This evidence must be seen in the context of the fact that aorta has both longitudinal and circular smooth muscle and adrenergic stimulation causes constriction of both types of muscle^11,19^. On the other hand, for the entire range of reduction in compliance, there was no change in mean aortic pressure, but the pulse pressure increased with reducing compliance almost linearly.

#### Simulation of small artery experiments (Figure 7)

R2 and C1 were initially set to values corresponding to calculations made from length and radius of one small artery sample as seen in Table 1.

##### Simulation 4

Increase in R2 alone by as little as 12% reduction in radius caused all 4 pressure parameters, namely, peak, trough, mean and pulse pressure to increase (top panels of Figure 7).

**Figure 7:**
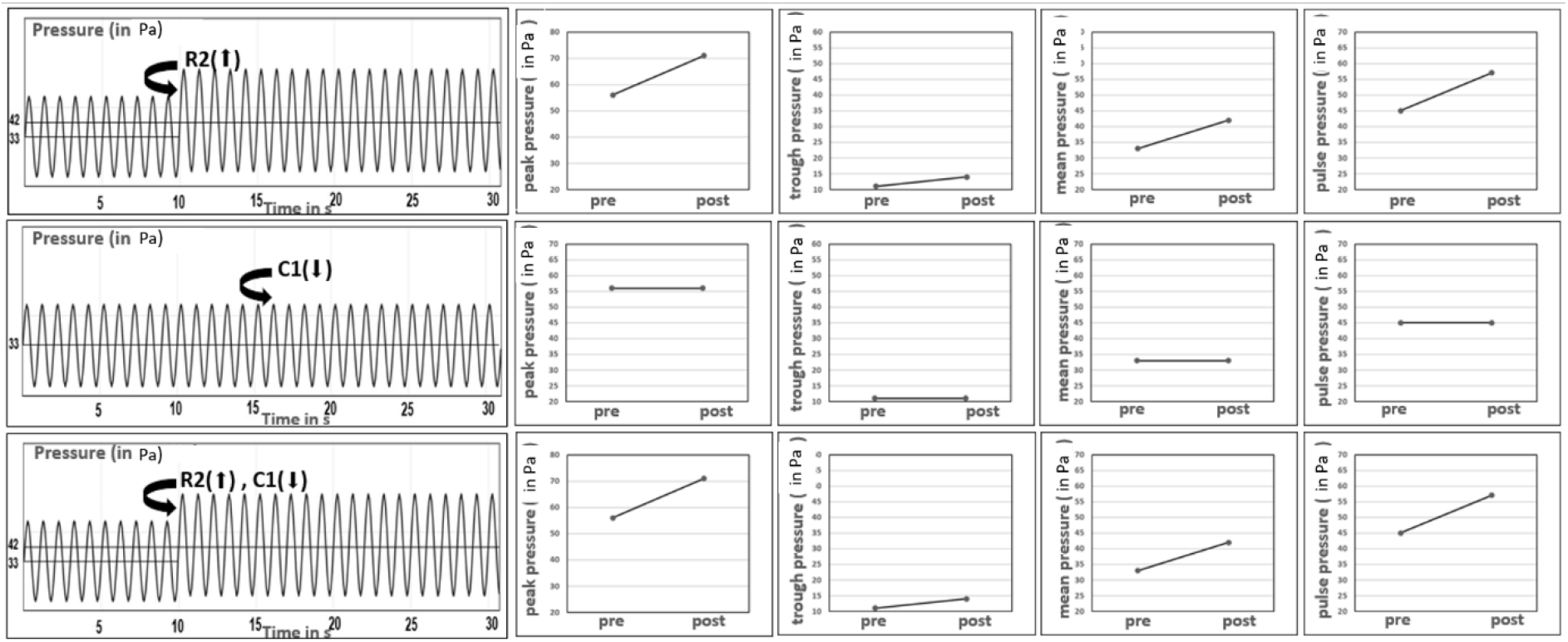
Simulations of small artery; increasing R2 (top panels); decreasing C1 (middle row of panels); increasing R2 and decreasing C1 simultaneously (bottom panels). V1 was set to 45Pa, 1Hz.

##### Simulation 5

Decrease in C1 by as much as 90% did not lead to any change in the 4 parameters (middle panels of Figure 7). This is probably because the value of C1 in small arteries is so low that any further decrease does not induce a change.

##### Simulation 6

Effect of combined changes in R2 (increase by 12% reduction in radius)) and C1 (decrease by 90% reduction in compliance) was the same as the effect of just increasing R2 (bottom panels of Figure 7).

Extending the calculations to cover all possible reductions in compliance, it was seen that any change in compliance does not lead to a change in mean pressure or pulse pressure (see, Figure 8, bottom panels).

**Figure 8:**
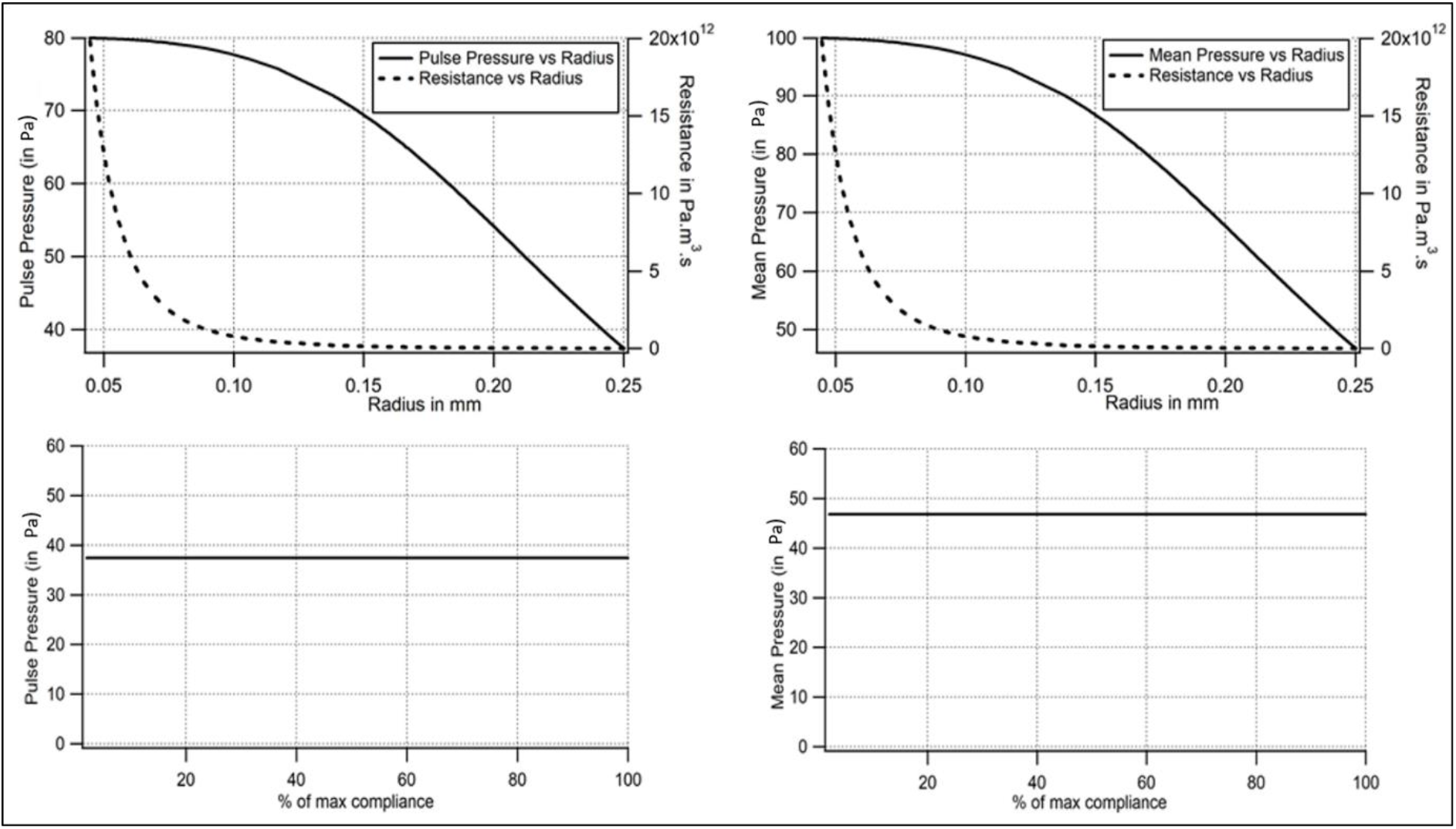
Effect of all possible reductions of radius on pulse pressure and mean pressure (top row of panels) in small artery simulations; Effect of all possible reductions in compliance on pulse pressure and mean pressure (bottom panels), in small artery simulations.

### Imaging experiments

Images of circular rings of aorta before and after addition of adrenaline were taken using an externally mounted camera and the images were analyzed using a calibrated custom software (n=3, Figure 9). Less than 20% reduction in radius from the initial value was observed in all three cases during vasoconstriction. Hence as discussed in Figure 6 simulation (top panel), a gross reduction in radius by more than 50%, that is essential to cause a significant rise in mean pressure, does not occur in aorta during vasoconstriction.

**Figure 9:**
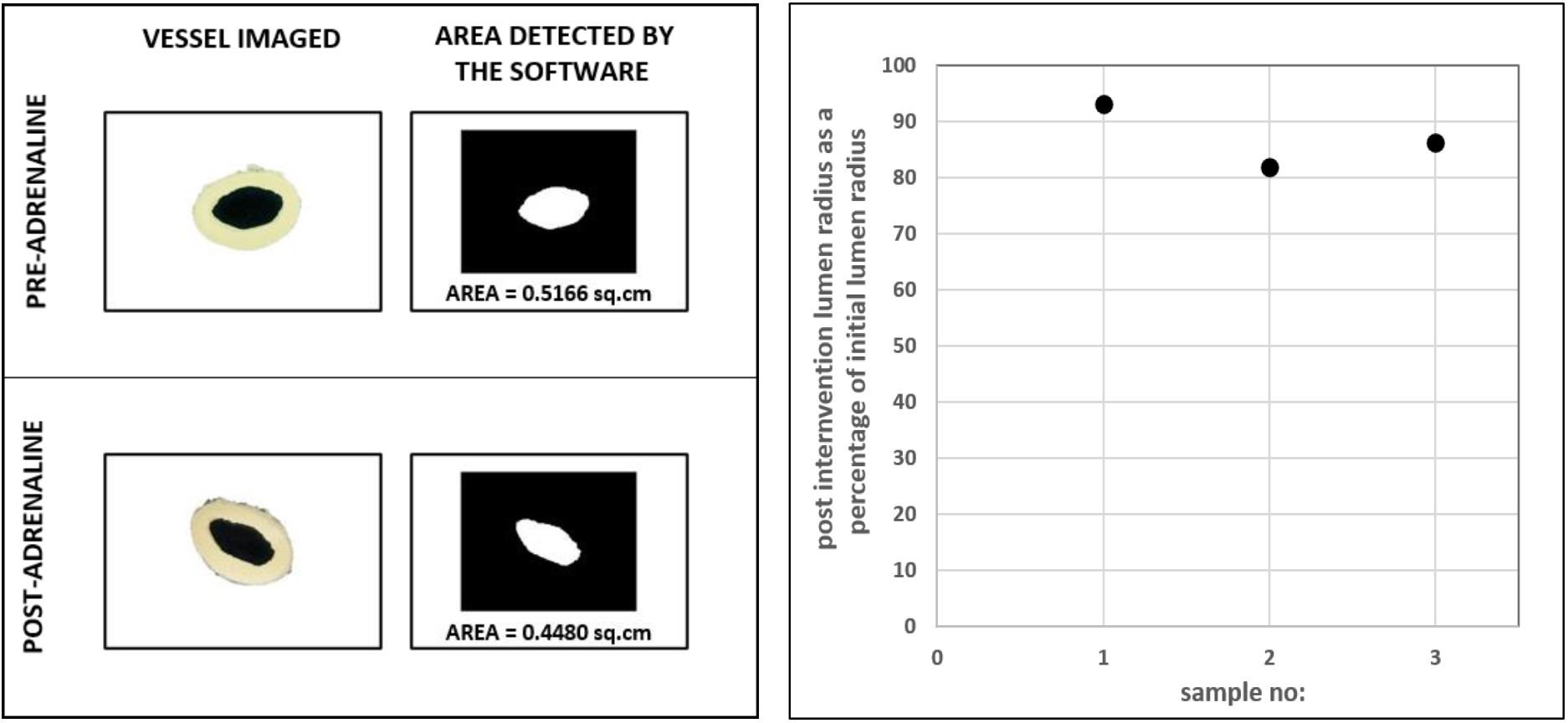
Imaging of circular rings of aorta pre and post-intervention

## Discussion

Generally, compliance in the vessels of systemic circulation is attributed to the aorta and large arteries. Compliance or its inverse elastance, of the aorta and large arteries is attributed largely to the presence of elastin fibres in the tunica media and the arrangement of collagen^20,21,22^. Due to wear and tear, the elastin fibres get damaged, the collagen mal-aligned, and such changes can lead to loss of elastance/compliance. This is suggested as the cause for isolated systolic hypertension (ISH) in older individuals^23,24,25^. If all the loss of compliance in age-related hypertension is due to loss of elastin, it must be irreversible with vasodilators which act by relaxing vascular smooth muscle. Missed out in a routine discussion of compliance in the context of hypertension is the contribution to it by the smooth muscle in the walls of large arteries. There is a considerable quantum of smooth muscle in the tunica media of large arteries and aorta. This smooth muscle is also arranged concentrically as in the arterioles. Constriction of smooth muscle in large arteries must lead to a decrease in compliance of the arteries. Such loss of compliance must be reversible, unlike the age-related loss of elasticity. In fact, it is reported that aortic compliance was lower in hypertensives than in normotensives and improved with nitroprusside infusion. The authors attribute the low compliance to vasoconstriction because of its reversibility^13^. Our experiments also demonstrate that the aortic compliance reduces with adrenaline.

The aortic segment will have a defined resistance as well in addition to compliance. The question we asked was if the aortic resistance increases with adrenaline. The response of the aortic segment to adrenaline, i.e., an increase in peak pressure, decrease in trough pressure and lack of change in mean aortic pressure suggests that the vasoconstriction caused by adrenaline decreases compliance but does not alter resistance of the aorta sufficiently enough to cause an increase in mean aortic pressure. This result is being categorically reported for the first time up to our knowledge.

Small arteries also have circular and longitudinal smooth muscle^11^. They too must have a defined resistance and compliance. Increase in mean pressure after addition of adrenaline is the expected response in these vessels, as these are resistance vessels and sympathetic stimulation is known to increase resistance. The increases in peak, trough and mean pressures can all be explained by an increase in resistance alone (due to a reduction in lumen diameter, resulting from contraction of circular smooth muscle)^26^.

Effect of adrenaline on small arteries in the biological experiments reported in Figure 4 resembles simulations 4 and 6. The observation with simulation 4 is in line with the known effect of adrenaline in increasing small artery and arteriolar resistance by reducing lumen diameter. Whether adrenaline reduces compliance of small arteries or not is a question that cannot be answered with these simulations, as a decrease in compliance did not lead to any change in pressure parameters. This can be attributed to the extremely low compliance of small arteries.

Considering the effect of adrenaline *in vivo*, the following may be inferred. The simulation experiments (Figure 6) show that, in the aorta, for an appreciable increase in resistance and therefore mean arterial pressure, the radius should reduce by 70%. As such gross vasoconstriction does not occur in the aorta *in vivo* (also confirmed in the imaging experiments reported in Figure 9), it may be concluded that in the intact animal, the increase in mean arterial pressure in response to vasoconstrictors is not due to vasoconstriction in the aorta (and large arteries). It is entirely due to vasoconstriction in the small arteries and arterioles.

The significance of the study is that in *in vivo* recordings, if resistance and compliance of the vasculature can be resolved on a beat-to-beat basis, using pressure and flow waveforms upon administering a drug, then any change in resistance can be construed as an effect on the small arteries and arterioles, and any change in compliance, as due to an effect on large arteries and aorta. Given that every drug profile may differ in terms of their site of action, this study fulfils the need of re-investigating the effect of every vasoactive drug on resistance vessels versus compliance vessels and thereby tailor treatment to correct the component at fault. For instance, administering vasodilators that further lowers diastolic pressure would be a contraindication to the treatment of ISH. Instead, a drug which causes vasodilation of large arteries and not the resistance vessels, would be ideal.

## Additional Information Section

### Data Availability Statement

All the data analyzed during this study are included in this published article. The datasets and simulation codes are available from the corresponding author on reasonable request.

### Author’s Translational Perspective

There is no definitive evidence upto our knowledge, if constriction of smooth muscles in large arteries contributes to resistance. We show here that there is no change in resistance or mean pressure during large artery vasoconstriction although compliance decreases. Whereas small artery vasoconstriction leads to an increase in resistance and hence an increase in mean pressure. This understanding would help redefine the actions of vasoactive agents currently used in the management of shock, ISH etc. For instance, administering vasodilators that further lowers the diastolic pressures would be a contraindication to the treatment of ISH. Instead, a reversal of the decreased compliance of large arteries would be ideal. Hence, it is crucial to understand if a drug acts at the level of small arteries or large arteries. Based on the results from this study, beat to beat resistance and compliance computed from intra-arterial pressure and flow measurements) under the effect of a drug can be resolved as differential responses of small and large arteries to vasomodulators, respectively. This would enable re-investigating the effect of every vasoactive drug on resistance vessels versus compliance vessels and thereby tailor the treatment to correct the component at fault. This would ensure targeted anti-hypertensive therapy based on one’s vascular biology.

### Author contributions

**NG**: Conceptualization, Data Curation, Formal analysis, Investigation, Methodology, Resources - instrumentation, experiments and analysis, Software, Validation, Visualization, Writing - Original Draft, Writing - Review & Editing. **AV** contributed to Methodology, Investigation, Validation, Writing - Review & Editing. **BJ** contributed to Conceptualization, Validation, Writing - Review & Editing. **SMZ** contributed to Investigation, Validation, Writing - Review & Editing. **SD** contributed to Conceptualization, Data Curation, Formal analysis, Methodology, Validation, Visualization, Supervision, Writing - Review & Editing. **GSK** contributed to Formal analysis, Resources, Validation, Visualization, Supervision, Writing - Review & Editing. **SS**: Conceptualization, Data Curation, Formal analysis, Investigation, Methodology, Resources –experiments and analysis, Validation, Visualization, Fund acquisition, Project Administration, Supervision, Writing - Original Draft, Writing - Review & Editing.

## Acknowledgements

The Authors thank the Department of Biotechnology (DBT), Government of India, for funding the study. The Authors are grateful to the Animal Ethics Committee, Christian Medical College, Vellore for approving the study. NG thanks Indian Institute of Technology Madras for fellowship support.

